# Recent advances in the study of fine-scale population structure in humans

**DOI:** 10.1101/073221

**Authors:** John Novembre, Benjamin M. Peter

## Abstract

Empowered by modern genotyping and large samples, population structure can be accurately described and quantified even when it only explains a fraction of a percent of total genetic variance. This is especially relevant and interesting for humans, where fine-scale population structure can both confound disease-mapping studies and reveal the history of migration and divergence that shaped our species’ diversity. Here we review notable recent advances in the detection, use, and understanding of population structure. Our work addresses multiple areas where substantial progress is being made: improved statistics and models for better capturing differentiation, admixture, and the spatial distribution of variation; computational speed-ups that allow methods to scale to modern data; and advances in haplotypic modeling that have wide ranging consequences for the analysis of population structure. We conclude by outlining four important open challenges: The limitations of discrete population models, uncertainty in individual origins, the incorporation of both fine-scale structure and ancient DNA in parametric models, and the development of efficient computational tools, particularly for haplotype-based methods.

---

If one assumes humans across the globe are a single randomly mating population, it would result in only a 5-15% average error when predicting the proportion of observed heterozygotes at a locus. This closeness to an idealized randomly mating population is one vestige of how little evolutionary time has passed since the common origin of all humans in Africa. The departure from random mating predictions due to population differentiation has a classic quantitative measure, called F_ST_, which appropriately takes on values of 5-15% in global samples of human populations [1–3]. If one zooms in within continental regions of the globe, F_ST_ tends to be even lower, regularly taking values below 1%, a threshold which we use here informally to define “fine-scale structure”. A triumph of the large datasets available to contemporary population geneticists is that they allow fine-scale structure to be detected and dissected to reveal population relationships (Figure 1).

**Figure 1:**
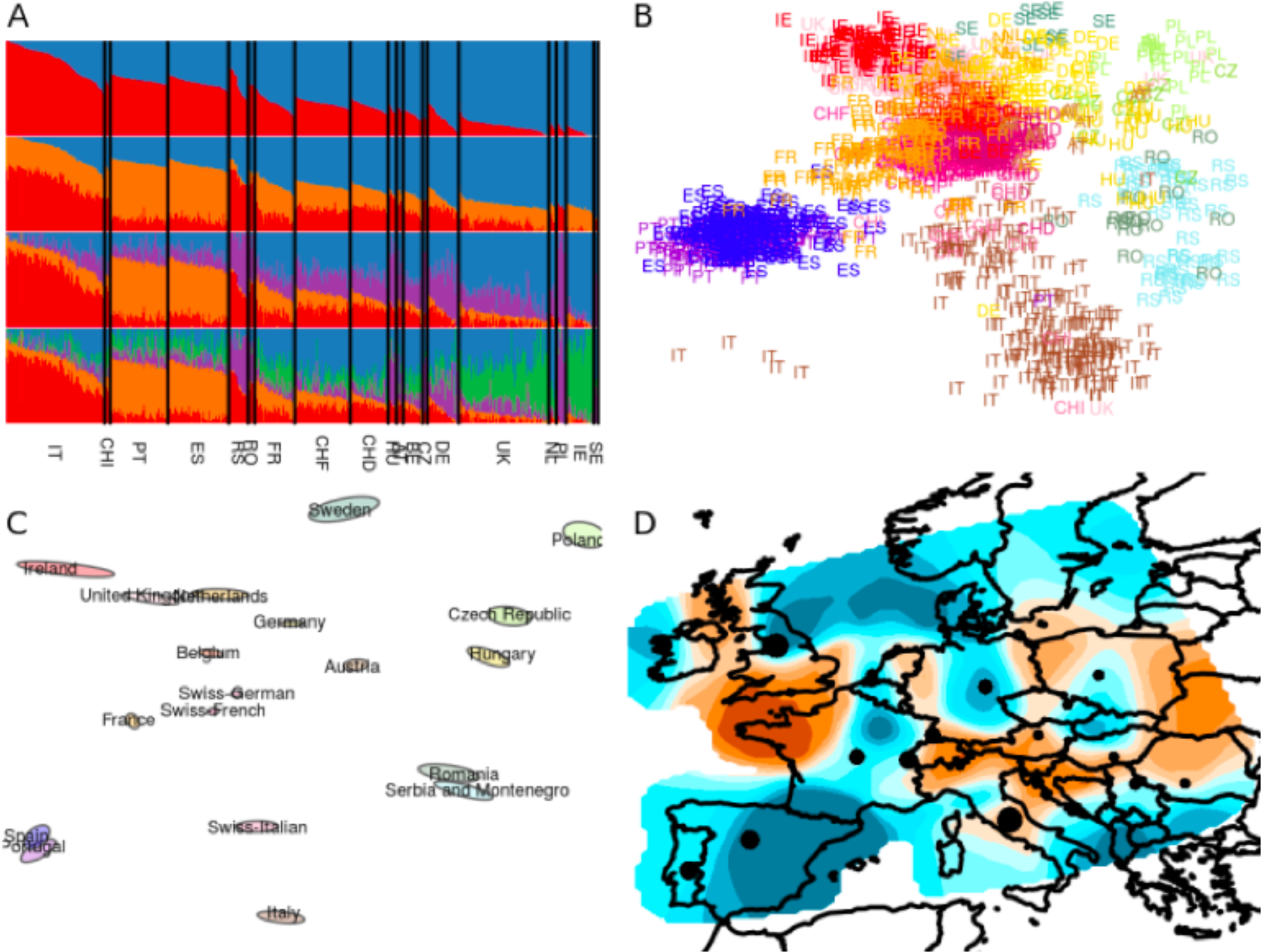
Four methods for assessing population structure using large-scale single-nucleotide polymorphism data. A) Ancestry proportion inference using ADMIXTURE. B) Principal components analysis. C) Plot of population in ‘geogenetic’ space using SpaceMix. D) Visualization of effective migration rates using EEMS (brown = low effective migration; blue = high effective migration). Each method is applied to the dataset analysed in [92] after filtering out populations with fewer than 10 individuals, where population identifiers are defined on the basis of grandparental ancestry. Structure is visible, even though F_ST_-values average 0.004 between broad geographic regions in Europe [92].

The reason large datasets allow fine-scale structure to be dissected is a statistical phenomenon [4]. Subtle differences in average pairwise similarity become more apparent as the scale of a dataset increases (Figure 2). For principal component analysis (PCA), Patterson et al. [5] have argued that structure reveals itself much like a phase change in physics; namely if the product of the number of genetic markers (m) and individuals (n) is greater than 1/(F_ST_^2^) then structure will be evident. The great fortune of human geneticists is that novel technologies have made it affordable to amass datasets large enough to characterize even very subtle population structure.

**Figure 2:**
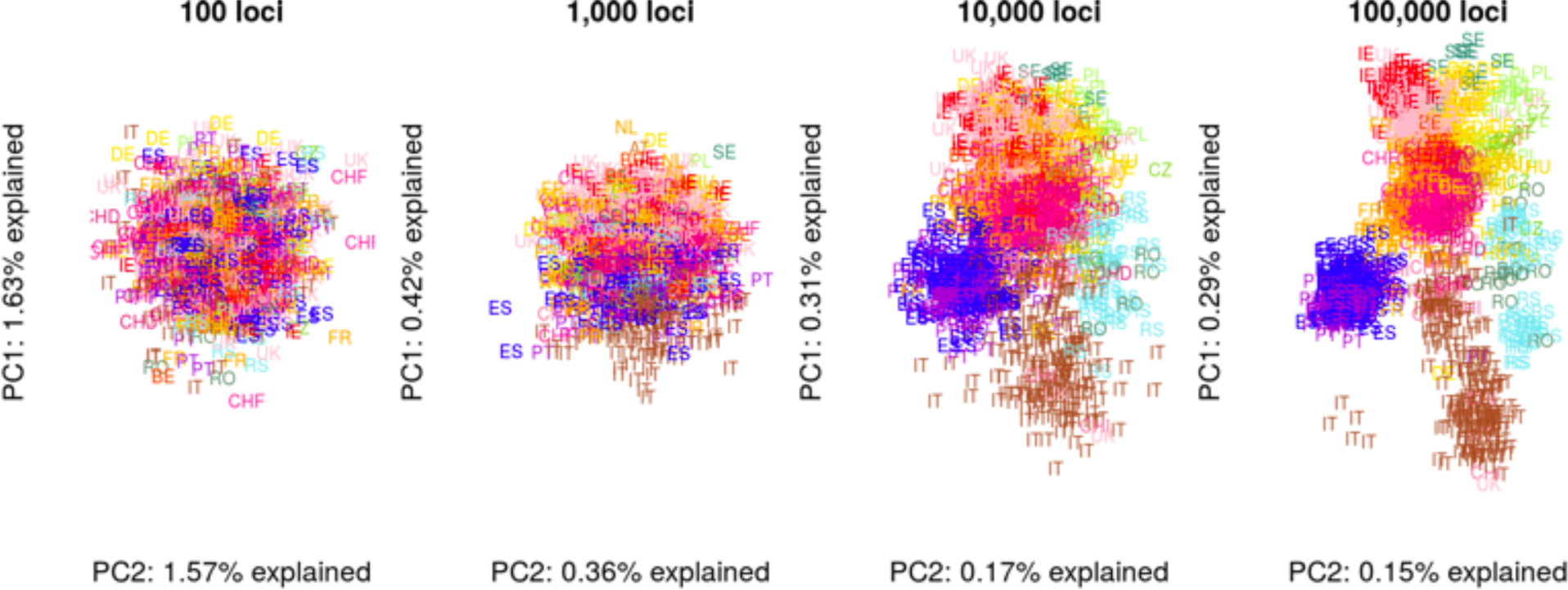
A large number of loci is required to reveal fine-scale population structure using PCA. Four subsamples with an increasing number of loci were taken from the[93] dataset. Using 100 loci, Europe appears panmictic, whereas 1,000 loci are sufficient to establish a North-South cline. With 10,000 and 100,000 loci, fine-scale details are revealed.

Characterizations of fine-scale population structure are increasingly empowering the study of human population history, helping genetics play a role integral with linguistics, archaeology, and history in the study of the human past, as first envisioned by Cavalli-Sforza and colleagues [6]. Studies of population structure have also allowed the medical genetics community to build more robust genome-wide association studies for disease risk [7–9], enabled evolutionary geneticists to identify exceptional regions in the genome that have undergone local adaptation [10–12], and facilitated the individual-level ancestry assessment that is increasingly popular in personalized genomics [13], though not without criticism [14].

The most obvious advances in the study of fine-scale structure are the result of increased genome-wide single-nucleotide polymorphism (SNP) and sequencing data being collected from diverse regions across the world [15–17]. More profoundly, the recent availability of genome-wide data from archaeological samples of modern humans (“ancient DNA”, aDNA) is revolutionizing our understanding of the processes that generated present-day population structure [18]. As one stimulating example, aDNA studies suggest that levels of differentiation in Europe may have actually decreased during the late Neolithic period and Bronze Age, as at least three major proposed population lineages have intersected through time [17,19,20].

In this review, we highlight several of the most exciting advances in analysis methods in the past three years. Analysis methods are especially crucial when structure is fine-scale, and as we show, there has been extensive progress in numerous directions. To complete a picture of recent studies of human population structure, we recommend several other recent reviews [21–27]. We also highlight the Peopling of the British Isles Project [28] and two recent ancient DNA papers [19,20] as examples from the vanguard of fine-scale structure analysis using modern and ancient human data.

## Developing and understanding metrics for population differentiation

While Wright’s F_ST_ has been a workhorse for describing population structure for decades, a novel set of f-statistics have become highly influential since their original publication [29,30] because of their utility in studying the recent admixture that is common in human history. In the past several years, the use of f-statistics has continued to develop; for example, Raghavan et al. [31] developed an ‘outgroup-f_3_’ statistic to provide a measure of similarity that is insensitive to population-specific drift, making it more interpretable than F_ST_ in many settings. The advent of sequencing data has also made possible a metric that is especially sensitive to recent population structure: the estimated time to the most recent common ancestor of shared doubleton variants [32]. Doubleton-based metrics are already showing their utility to detect fine-scale structure in several large sequencing studies [33–35].

Alongside the expansion of new metrics, there is an increasing understanding of the basic properties of existing ones. One area of concern is that F_ST_ has inconsistent values across different types of loci (e.g. microsatellite, SNP array, and sequence data differ by 0.05-0.07 in several meaningful human examples [36,37] also see [38]). As demonstrated by Jakobsson et al. [37], this discrepancy is largely explained by the frequency of the most frequent allele at a locus, which differs greatly between marker classes [37]. However, the statistical strategy used also has an impact, in particular when rare alleles are present [36]. Similar understandings need to be developed for f-statistics. In that vein, one of us [39] recently showed how f-statistics can be understood in general coalescent terms, with expectations that can be derived under arbitrary parametric population history models. Interestingly, the work revealed that the mean number of pairwise differences between populations is a better measure of differentiation than the outgroup-f3 statistic.

## Refining and expanding models that handle human admixture

Recognizing that simple bifurcating trees are a poor model for human genetic diversity, several researchers have developed tree-building methods that take admixture and migration into account [29,40–42]. These methods typically build a guide tree and then add migration edges to represent recent admixture events. These methods represent a substantial advance and they are being used broadly; however, challenges remain with identifiability, exploring the space of all possible graphs, robustly selecting the number of migration edges, and exploiting tools from similar approaches developed independently in phylogenetics [43,44].

A more longstanding approach to study admixture is the use of global [45] and local ancestry models [46], and both have been advanced recently by computational speed-ups. When genome-wide SNP data first became available, difficulties in applying the now classic Bayesian method for admixture inference (STRUCTURE [45]) occasioned the development of fast maximum-likelihood based approaches [47,48]. The past two years have seen the return of Bayesian approaches with two new fast approaches that use variational approximations [49,50]. Impressively, teraSTRUCTURE [50] handles samples containing millions of individuals. Local ancestry approaches have recently been improved by the development of fast algorithms that leverage rare variants within ancestries [51] and wavelet techniques [52]. Unfortunately, distinguishing local ancestries among weakly differentiated source populations remains difficult. For these cases, alternative approaches based on admixture linkage disequilibrium, which sidestep local ancestry inference, have proven fruitful for detecting admixture occurring in the recent past (e.g. within the last three thousand years) and suggest that admixture is widespread in human populations [53–55].

Latent factor models, such as sparse factor analysis and PCA, are also applicable for inference of admixture [56,57]. For PCA, more efficient algorithms have been implemented, allowing application to hundreds of thousands of individuals with reasonable run-times [58,59]. Another useful development has been the expansion of Procrustes approaches (used to align PCA and geography data [60]) to allow merging of samples with low-coverage sequence data into PCAs from heavily genotyped reference panels [61,62]. Novel factor models are also being developed that may have advantages for assessing admixed samples and controlling structure in association studies [63].

## Putting geography into studies of population structure

Geographic information has been under-utilized in the study of fine-scale structure despite its central importance in the process of generating structure. Several recent approaches make progress by using Wishart distribution to model genetic similarity as a function of spatial distance. In one approach, Bradburd et al. [64] use a covariance model to visualize samples in a “geogenetic space”. In homogenous isolation-by-distance scenarios, the geogenetic space should mirror the geographic space. Deviations from homogeneity will result in adjusted placement of populations in geogenetic space. For example, barriers to migration result in larger geogenetic distances between populations. In some respects the methods should behave like PCA on spatial data [65], but with less of the sensitivity to uneven sample sizes that is typical of PCA [57,65]. Discrete admixture events can be accounted for by adding admixture links between sources and targets, in a manner similar to how recent tree-based methods add migration edges to model admixture (see above).

A second approach using the Wishart distribution is the EEMS method [66] which uses observed pairwise genetic similarities to estimate a map or “surface” of effective migration rates. The inferred migration rates are “effective” in that they reflect migration proportions scaled by effective population sizes under an equilibrium model. As few empirical systems (and especially humans) are at equilibrium, the surfaces should best be interpreted as a tool for visualizing patterns of genetic differentiation relative to geographic distance. In closely related work, Hanks and Hooten [67] independently develop a Wishart framework much like that in EEMS that can test whether a particular environmental variable is predictive of migration rates. Two additional exciting advances, not using the Wishart distribution, are a method (localdiff [68]) that uses locally computed F_ST_ values to visualize barriers, and an approach that models F_ST_ as a function of the bearing between two populations to study anisotropic patterns of spatial differentiation [69].

## Extracting signatures of fine-scale structure in haplotype data

A very active and promising arena of research is in the development of haplotype-based methods for studying fine-scale structure. Local ancestry models (see above) have long been the only haplotype-based approach used to study structure, and haplotypes can be more informative for assignment [70,71], but a bevy of methods are being developed that leverage haplotypes in novel ways.

Methods using long shared haplotypes (also known as tracts of identity-by-descent, IBD) are particularly well-suited for the characterization and interpretation of fine-scale population structure [72]. As an example of their power, Ralph and Coop [73] showed that IBD patterns in Europe can reveal more subtle structure than simple pairwise SNP similarity. More recently Baharian et al [74] use the spatial distribution of long shared haplotypes to estimate dispersal rates in the context of the African-American history. These methods have exceptional promise, though interpretations can be complicated when shorter tracts are considered [75]. A related approach to IBD patterns is embodied in the fineSTRUCTURE model [76,77] which uses long shared haplotypes detected through the use of the Li and Stephens haplotype-copying model [78] and then processes them through a downstream analysis that includes mixture modeling of copying profiles. This model underlies the striking structure revealed in the Peopling of the British Isles project [28].

A second approach has been to focus on full chromosomal haplotype data using approximate coalescent models. Using the Sequential Markov Coalescent (SMC) approximation, it is now possible to study population divergence using small numbers of genomes [79,80]. As one example, Schiffels and Durbin suggest a novel non-parametric approach based on inference of cross-coalescent rates through time [80,81]. In related work, a recent approach for sampling coalescent genealogies genome-wide [82] is a remarkable achievement but has yet to be adapted to explicitly study population structure.

A drawback of most haplotype-based methods is that they are computationally very expensive. A major breakthrough is the development of fast and scalable algorithms for representing haplotype data in ways that make haplotype similarity evident and easy to query. One such algorithm, the Positional Burrows-Wheeler Transform (pBWT, [83]) is revolutionary in this regard, and a new extension [84] is also exciting as it links the Li and Stephens haplotype-copying model to the pBWT framework and provides approaches that can handle unphased data.

## Open challenges

Despite all the methodological progress, many fundamental challenges exist for fine-scale population structure studies. Many of these challenges stem from the difficulty of encapsulating the complexity of human population structure with mathematical models.

On the one hand, many models assume a small number of discrete, temporally continuous populations as the units of analysis. This is only an approximation to the structure of human populations on the ground, and there is a strong trade-off between the ease-of-analysis provided by using a small number of discrete populations and the error induced due to model violations. On the other hand, other models are not sufficiently parametric. Many available methods focus on summarizing the observed population structure in a form that facilitates interpretation, but without explicitly modeling the historical processes that shaped these patterns.

As a symptom of this problem, we currently lack a widely accepted generative model for human fine-scale data. Put another way, we do not have a clear simulation protocol to produce “realistic” human data with fine-scale structure. This hinders applications such as testing new methods and evaluating evolutionary models of disease variants in human populations. As an example, consider how recent aDNA studies have made clear that fine-scale structure in Europe is partly driven by temporally dynamic admixture patterns between 3 ancestral populations [17, 19, 20]; it is unclear from existing publications what pairing of simulation protocol and parameters would generate an *in silico* whole-genome dataset that replicates the basic features of European fine-scale structure.

A related persistent challenge is that choices must be made regarding assigning individual origins based on sampling location, individual birthplace, or an origin based on parental or grandparental ancestry. Given the ubiquity of human movement and admixture, the choice can complicate interpretations and/or result in samples being omitted when origins are unclear (e.g. grandparents of differing origins). The problem also arises when assigning aDNA samples into analysis units when they might vary by location, cultural context, and sampling time. One must always interpret results with the location assignment procedure in mind. Ideally, approaches can be developed that more explicitly model the uncertainty in this stage of analysis.

Another practical challenge with fine-scale population structure is that analysts must be especially cautious regarding model deviations such as ascertainment bias, heterogeneous linkage disequilibrium (LD) patterns, and complex mutational processes. For example, even after basic LD filtering, PCs can be affected by large blocks of SNPs with complex patterns of LD, such as those that arise due to structural variants like the polymorphic 4Mb Chr 8p23 inversion in Europe (e.g. PC2 in the study of [85]). Such patterns likely pollute the results of various methods that assume marker independence as well as haplotype-based methods that do not model complex LD patterns due to structural variants. One precaution is to inspect the PC loadings and repeat the PCA after removing any genomic regions with exceptionally high loadings. An example of a mutagenic process potentially influencing haplotype-based approaches is the error-prone DNA-Polymerase Zeta, which may introduce dinucleotide and other aberrant mutation patterns [86] that bias analyses when handled as distinct mutations.

A final precaution, and one of broader societal relevance, is that a viewer can become misled about the depth of population structure when casually inspecting visualizations using methods such as PCA, ADMIXTURE, EEMS or fineSTRUCTURE. For example, untrained eyes may overinterpret population clusters in a PCA plot as a signature of deep, absolute levels of differentiation with relevance for phenotypic differentiation. This is an ironic inverse of what Edwards harshly termed Lewontin’s fallacy [4], and what we might instead call Lewontin’s nightmare. To prevent these misinterpretations, first we encourage practitioners to make absolute metrics of differentiation clear to audiences (e.g. F_ST_, PCA proportion of variance explained). Weak levels of differentiation, as measured by F_ST_, imply that neutral quantitative traits will be weakly differentiated as well [3,87]. Second, visually displaying the geographic distribution of a manageable number of random markers from a dataset can be helpful for students and broader audiences to gain a direct sense of levels of population structure. Several resources make this feasible for human genetic datasets (GGV browser [88], ALFRED [89,90] and HGDP Selection Browser [91]).

Without any question, the study of fine-scale structure has been an exciting frontier of contemporary population genetics, with extensive progress and continued promise. As this work continues, we will begin to more fully understand the processes that shape fine-scale structure in humans, and have a more full perspective on human origins. Of broader relevance, this progress also provides guidance for studying other species with highly dynamic population histories, and many of the methods reviewed here are useful for applications outside of humans.

## Acknowledgements

J.N. would like to acknowledge NIH support (R01HG00708, GM108805, 1U01 CA198933-01), and B.P. would like to acknowledge a Swiss NSF Postdoctoral Fellowship. We would also like to thank the members of the Novembre lab as well as Choongwon Jeong, Sohini Ramachandran, and Noah Rosenberg for helpful comments and discussions. The POPRES data used in the figures was accessed via dbGAP study accession phs000145.

